# Failure to reactivate salient episodic information during indirect and direct tests of memory retrieval

**DOI:** 10.1101/189522

**Authors:** Mason H. Price, Jeffrey D. Johnson

## Abstract

Several fMRI and EEG studies have demonstrated that successful episodic retrieval is accompanied by the reactivation of cortical regions that were active during encoding. These findings are consistent with influential models of episodic memory that posit that conscious retrieval (recollection) relies on hippocampally-mediated cortical reinstatement. Evidence of reactivation corresponding to episodic information that is beyond conscious awareness at the time of memory retrieval, however, is limited. A recent exception is from an EEG study by Wimber, Maaβ, Staudigl, Richardson-Klavehn, and Hanslmayr (2012) in which words were encoded in the context of highly salient visual flicker entrainment and then presented at retrieval in the absence of any flicker. In that study, coherent (phase-locked) neural activity was observed at the corresponding entrained frequencies during retrieval, consistent with the notion that encoding representations were reactivated. Given the important implications of unconscious reactivation to past findings and the modeling literature, the current study set out to provide a direct replication of the previous study. Additionally, an attempt was made to extend such findings to intentional retrieval by acquiring EEG while subjects were explicitly asked to make memory judgments about the flicker frequency from encoding. Throughout a comprehensive set of analyses, the current study consistently failed to demonstrate evidence for unconscious reactivation, and instead provided support that test items were indistinguishable according to their prior encoding context. The findings thus establish an important boundary condition for the involvement of cortical reinstatement in episodic memory.

## 1. INTRODUCTION

Neuroimaging studies of episodic memory have demonstrated that brain regions active when an event is encoded are at times reactivated during its retrieval (for reviews, see Rugg et al., 2008; Danker & Anderson, 2010; Rissman & Wagner, 2012; Rugg et al., 2015). In the context of psychological theory, such findings align with the principle of transfer-appropriate processing, whereby retrieval success is more likely when cues are processed in a manner similar to the processing at encoding (Morris et al., 1977; also see Tulving & Thomson, 1973). Likewise, the notion of encoding-retrieval similarity is central to prominent neurobiological models of episodic memory. According to these models, cortical activity patterns corresponding to the neurocognitive processes and representations engaged at encoding are represented sparsely by the hippocampus (Marr, 1971; Teyler & DiScenna, 1986; Alvarez & Squire, 1994; McClelland et al., 1995; Rolls, 2000). Later presentation of an effective retrieval cue activates the hippocampal representation, which in turn gives rise to reinstatement of the original cortical pattern. Common to both the psychological and neurobiological accounts is the prediction that encoding-retrieval similarity will be particularly strong during instances of conscious recollection (Hasselmo & Wyble, 1997; Norman & O’Reilly, 2003; also see Norman, 2010), as evidenced by either greater likelihood of reported retrieval success or with access to additional details not present in the cue. The current study tests whether the phenomenon of cortical reinstatement also extends to situations in which salient information present during encoding is outside of conscious awareness at the time of retrieval.

To investigate cortical reinstatement, studies generally take an approach of identifying the similarity (or overlap) between neural correlates of encoding and retrieval in the context of different subjective measures of memory (e.g., Wheeler et al., 2000; Kahn et al., 2004; Johnson & Rugg, 2007). In one study using functional magnetic resonance imaging (fMRI), Johnson et al. (2009) employed multivariate pattern analysis (MVPA; see Haynes & Rees, 2006; Norman et al., 2006; Rissman & Wagner, 2012) to test for retrieval-related reactivation of cortical patterns corresponding to three elaborate encoding tasks. Reactivation was greatest in magnitude when subjects indicated that retrieval was accompanied by the recollection (“remembering”) of encoding details (also see Kahn et al., 2004; McDuff et al., 2009; Staresina et al., 2012), consistent with the proposal that hippocampally-mediated reinstatement is associated with conscious retrieval. However, the involvement of reinstatement in episodic retrieval was additionally expanded to include non-recollective memory judgments. In particular, when subjects were reportedly unable to recollect details from encoding, they rated their level of confidence (in “knowing”) that an item was old or new. Reactivation was evident when old judgments were associated with high confidence, thus challenging the notion that reinstatement is restricted to instances of recollection. These findings instead suggested that cortical reinstatement might reflect a continuous neural signal that is informative to the subjective characteristics of memory retrieval (Mickes et al., 2009; Wixted & Mickes, 2010).

The “remember/know” procedures highlighted above, as well as source memory tasks that require an overt response to specific information from encoding (e.g., McDuff et al., 2009; Kuhl et al., 2011; for EEG evidence, see Waldhauser et al., 2016), encourage subjects to attempt to retrieve episodic details intentionally. The interpretation of any reactivation effects from these procedures is thus inconclusive as to whether they are elicited by the retrieval cue or generated by the subject in preparation for the upcoming item. A recent study by Kuhl et al. (2013), however, used a modified source memory task to further extend the role of cortical reinstatement beyond intentional retrieval and into the domain of incidental retrieval. Subjects in that study encoded words paired with either a picture of a face or a scene, and picture presentation was additionally manipulated to be to either side of a central fixation. Memory tests then used the words as cues to probe either the category of the paired picture (face/scene) or its location (left/right). MVPA revealed that category-related reactivation evident in several frontal and parietal cortical regions was stronger when the category compared to the location was probed, consistent with the notion that cortical reinstatement can be modulated by the intention (or goal) to recollect. By contrast, reactivation in the medial temporal lobe was equally strong for the two test types. The latter finding suggested that, even when the retrieval demands explicitly focused on location information from encoding, reinstatement corresponding to incidental retrieval of the salient, but task-irrelevant, category information still occurred.

Although fMRI studies such as those described above have broadened the functional significance of reinstatement in episodic memory, they fall short in potentially identifying another limiting condition of cortical reinstatement: whether it can occur for information that is outside of conscious awareness. Despite the prominent focus on reinstatement in providing conscious access to additional details at retrieval (Hasselmo & Wyble, 1997; Norman & O’Reilly, 2003), the ability of the hippocampus to function in an obligatory manner for both encoding representations and re-engaging them upon presentation of an effective cue has also been proposed (Moscovitch, 1992, 2008). However, the slow temporal resolution of fMRI leaves open the ambiguity that any observed effects could reflect either slow, and likely conscious, reactivation or rapid, and perhaps unconscious, reactivation (for related discussion, see Kuhl et al., 2013). A recent study by Wimber et al. (2012) alternatively used electroencephalography (EEG) to address the possibility of unconscious reactivation during memory retrieval. Subjects encoded a series of words presented in the context of visual frequency entrainment, whereby the background flickered at one of two frequencies (6 and 10 Hz). On a later recognition test, words were shown in the absence of the flicker. The main reported finding was that word retrieval was accompanied by phase-locked activity at the corresponding frequency of the prior flicker, consistent with the notion that the encoding information was reactivated. Importantly, the frequency-related effects were evident within about 300 ms after onset of the test words, and a separate behavioral experiment indicated that subjects were at chance when explicitly asked to identify the flicker rate previously associated with each word. Together, these findings suggest that reactivation can occur not only in an obligatory (automatic) manner with presentation of the retrieval cue, but outside of conscious awareness.

Given the novelty of the aforementioned results and their challenge to the widely held position that cortical reinstatement is associated with conscious retrieval, the current study sought to provide a direct replication and extension of those findings. The behavioral procedures were designed to follow closely those of Wimber et al. (2012), including the nature and timing of the stimuli, the tasks performed, and the number of items comprising each phase. EEG data were continuously acquired throughout the encoding and memory retrieval phases. At encoding, subjects completed a syllable-judgment task for a series of words, while the background flickered at either 6 or 10 Hz. After a short delay, a recognition memory test was administered in which subjects judged their confidence about the old/new status of each word in the absence of any flicker. As this test made no explicit reference to retrieving the prior flicker condition, it is hereafter referred to as the *indirect test*.^1^ In a subsequent *direct test* phase, EEG data was acquired while the same subjects were asked about the flicker condition associated with each word presented at encoding. Compared to the indirect test, the direct test allowed for examination of cortical reinstatement under more ideal conditions when the flicker information was subject to intentional retrieval (e.g., Kahn et al., 2004; Johnson & Rugg, 2007; Johnson et al., 2009; McDuff et al., 2009; Kuhl et al., 2011; Waldhauser et al., 2016).

## 2. RESULTS

All of the analysis scripts used to generate the results and figures reported here, along with the relevant behavioral data and processed EEG data, are provided on the Open Science Framework (OSF) at https://osf.io/86wka/. The raw and transformed EEG data are not included, due to space constraints, but are available from the authors on request.

### 2.1. Behavioral results

During the encoding phase, response times (RTs) were statistically equivalent (t_17_ = 1.10, p = .29, BF_01_ = 2.42) for words presented with 6-Hz (M = 2694, SD = 654) and 10-Hz visual flicker (M = 2687, SD = 648).

The mean response proportions from the indirect test are provided in Table 1. Collapsing across confidence, the overall hit rates were equivalent for words from the 6- and 10-Hz encoding conditions (M = .74 and SD = .13 for each condition; t_17_ = .49, p = .63), as were the associated RTs (respectively, M = 2316 and 2305 ms, both SD = 529; t_17_ = .94, p = .36). Bayes-factor (BF) t-tests of these data revealed moderate support for the null hypothesis in both cases (BF_01_ = 3.70 and 2.79 for hit rates and RTs, respectively; Rouder et al., 2009). A two-way ANOVA of the response proportions for old words, with factors accounting for confidence (all 6 levels) and prior frequency condition (6-vs. 10-Hz), gave rise to a significant main effect of confidence (F_1.3,21.5_ = 38.26, p < .001) but no effects involving the frequency factor (Fs < 1). (Degrees of freedom for this and subsequent analyses were Greenhouse-Geisser adjusted when appropriate.) An analogous BF ANOVA indicated that a model with only the confidence main effect (BF_10_ = 1.76 × 10^47^) was preferred over a model with both main effects by a factor of 6.70 and over a full model by a factor of 240.82 (Rouder et al., 2012). Because old words were typically designated with the “sure old” response, it was not possible to compute meaningful statistics on the RTs at each of the remaining confidence levels. No significant difference was observed, however, between “sure old” RTs for words from the 6-Hz (M = 2264 ms, SD = 553) and 10-Hz (M = 2249 ms, SD = 561) conditions (t_17_ = 1.16, p = .26; BF_01_ = 2.30).

**Table 1:** Mean (SD) proportion of each response type during the indirect test.

| Trial type | Response type |  |  |  |  |  |
| --- | --- | --- | --- | --- | --- | --- |
|  | sure old | probably old | maybe old | maybe new | probably new | sure new |
| old, 6 Hz | .51 (.23) | .15 (.07) | .08 (.08) | .05 (.05) | .11 (.06) | .07 (.04) |
| old, 10 Hz | .50 (.22) | .15 (.07) | .09 (.08) | .07 (.06) | .11 (.05) | .07 (.05) |
| new | .12 (.10) | .10 (.06) | .07 (.06) | .12 (.07) | .21 (.10) | .35 (.20) |

For the direct test, the overall rate of correctly identifying the prior frequency condition was .49 (SD = .03) and not significantly different from chance (.50; t_15_ = .77, p = .45; BF_01_ = 3.02). Near-chance performance was also evident at the level of individual subjects, with the highest correct response rate being .53. The data separated according to the prior flicker conditions indicated that correct response rates were slightly higher, but not significantly so (t_15_ = 1.83, p = .09; BF_01_ = 1.01), for words from the 10-Hz (M = .52, SD = .08) compared to 6-Hz condition (M = .47, SD = .06). This difference could be interpreted as a slight bias to respond “fast” (“10-Hz”), but a computed bias index (B_r_, based on arbitrarily treating 10- and 6-Hz trials as old and new, respectively; Snodgrass & Corwin, 1988) did not differ from neutral (M = .52, SD = .06; t_15_ = 1.40, p = .18; BF_01_ = 1.72). For 6-Hz trials, the mean RTs for “slow” and “fast” responses were 2268 (SD = 519) and 2237 ms (SD = 526), respectively; for 10-Hz trials, the respective mean RTs were 2269 (SD = 511) and 2232 ms (SD = 522). A two-way ANOVA of these data gave rise only a significant effect of response type (F_1,15_ = 5.72, p < .03; Fs < 1 for the other effects), reflecting the impression that “fast” responses were faster than “slow” responses. A BF ANOVA indicated support for the model comprising only the response type main effect (BF_10_ = 2.22), which was preferred over the model with both main effects by a BF of 3.93 and over the full model by a BF of 12.15.

### 2.2. EEG results

Analyses of the EEG data first focused on assessing the inter-trial coherence (ITC) associated with the visual flicker at encoding and then applying an analogous approach to data from the indirect and direct tests. As described in the Introduction, ITC differences during the indirect test according to the prior flicker frequency would constitute evidence for the reactivation of encoding information. By combining such evidence with the behavioral results from the direct test, whereby subjects were unable to accurately identify the prior frequency condition of words (see above), the reactivation could be labelled as occurring incidentally (and perhaps unconsciously). Additionally, because subjects were explicitly asked during the direct test to retrieve the frequency information associated with each word, ITC differences in that phase were expected to provide a stronger test for reactivation.

#### 2.2.1. Encoding phase

ITC was first contrasted across the two visual flicker conditions of the encoding phase. Due to the constant nature of the flicker, ITC differences were expected to be evident throughout the stimulation period (0–2000 ms). As shown in Figure 1A, sustained ITC differences between 6- and 10-Hz trials were present across the frequency spectrum for the data collapsed over all electrodes. Notably, ITC was higher at 6 Hz for 6-Hz compared to 10-Hz trials, and at 10 Hz for 10-Hz trials. To test the effects in these two frequency bands of interest, the ITC differences were collapsed over the stimulation period by averaging the data from 5.5–6.5 Hz and 9.5–10.5 Hz for the respective bands (see Wimber et al., 2012, for similar use of band ranges). The difference in each band reached significance (6 Hz: t_17_ = 8.21, p < .001; BF_10_ = 61864; 10 Hz: t_17_ = 5.45, p < .001; BF_10_ = 598.33). To further probe the reliability of these effects, ITC differences were separately tested at each time point and frequency band, the results of which are displayed in Figure 1B. The effects in both frequency bands of interest passed the cluster-corrected threshold for significance (6 Hz: k = 486, p < .003; 10 Hz: k = 500, p < .001). Additional effects were evident at higher (and sometimes harmonic) frequencies, such that ITC was higher for 6-Hz trials at about 15 Hz (k = 329, p < .017) and for 10-Hz trials at about 20 (k = 738, p < .001) and 30 Hz (k = 312, p < .016).

**Figure 1.**
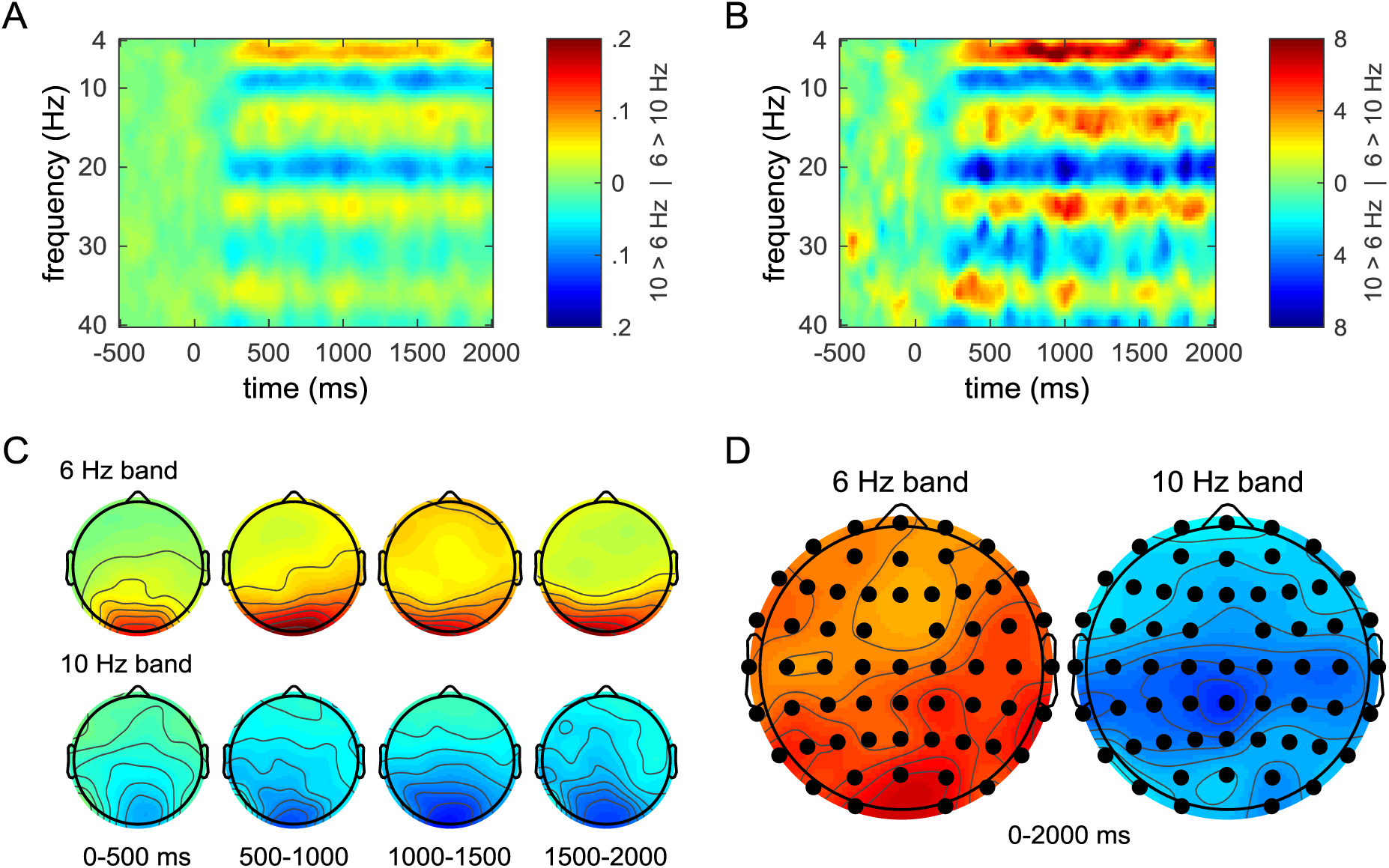
Inter-trial coherence (ITC) during the encoding phase. (A) Group-averaged time-frequency plots of ITC differences, collapsed over all electrodes. Warm colors indicate higher ITC for 6-Hz trials; cool colors indicate higher ITC for 10-Hz trials. (B) T-statistics associated with the differences shown in panel A, with warm and cool colors respectively indicating higher ITC for 6- and 10-Hz trials. (C) Topographic maps of ITC differences for the 6-Hz (top row) and 10-Hz (bottom row) frequency bands, averaged into consecutive 500-ms latency periods. The color scale from panel A is used. (D) Topographic maps of t-statistics for differences in the 6- and 10-Hz frequency bands across the 0–2000 ms period. The color scale from panel B is used. Significant differences (p < .05, one-tailed) were evident at every electrode, as indicated by the closed black circles.

The ITC differences at encoding were also expected to be maximal over the posterior scalp, given the visual modality of the flicker. Figure 1C displays the topographic maps of the effects at 6 and 10 Hz (again averaged over 5.5–6.5 Hz and 9.5–10.5 Hz ranges), collapsed into 500-ms intervals for descriptive purpose. As expected, the differences maintained a posterior maximum across the intervals. Collapsing over the entire stimulation period (0–2000 ms) revealed maximal differences at the O1 electrode for the 6-Hz band (M = .18, SD = .12, t_17_ = 6.33, p < .001) and at the POz electrode for the 10-Hz band (M = .13, SD = .17, t_17_ = 2.99, p < .01). Moreover, statistically testing the ITC differences for the entire period at each electrode, as shown in Figure 1D, gave rise to effects in each band that were in the predicted direction and significant at every electrode. The electrode-wise t-values ranged from 3.22 to 6.69 for the 6-Hz band, and from 2.24 to 5.88 for the 10-Hz band. Unsurprisingly, these effects surpassed the corrected threshold for number of clustered significant electrodes (p < .001 for each band). (For the 6- and 10-Hz bands, respectively, clusters of 17 and 19 electrodes corresponded to the critical threshold of p < .05.)

#### 2.2.2. Indirect test

Having established ITC effects at encoding, EEG data from the indirect test were next analyzed for analogous differences indicative of reactivating frequency-related information. Figure 2A shows the ITC differences across the recording epoch, collapsed across all electrodes, for old words eliciting correct responses (i.e. hits, regardless of confidence). (Similar results were obtained when using the hits associated with only “sure old” responses, as reported in the Supplemental Material.) The corresponding t-values from statistically contrasting ITC at each frequency band and time point are shown in Figure 2B, but no differences surpassed the cluster-corrected significance threshold (k > 110 and 111 corresponded to one-tailed p < .025 for 6-Hz and 10-Hz trials, respectively). The largest cluster exhibiting higher ITC for 6-Hz trials was 32 data points in size (p = .813), whereas the largest cluster of the opposite effect was 40 data points (p = .644). Since reactivation effects might be expected to occur with word onset and be short-lived, ITC differences collapsed across the 0–300 ms period were also analyzed for the 6- and 10-Hz bands (also see Wimber et al., 2012). However, there was no evidence of reactivation during this period (at 6 Hz: t_17_ = .72, p = .48; at 10 Hz: t_17_ = .78, p = .45) and instead, there was moderate evidence for the null effects (BF_01_ = 3.26 and 3.15, respectively). A longer period of 0–500 ms, given that it still preceded the typical left parietal old-new effect (see Rugg & Curran, 2007, for review; also see the Supplemental Material), was also analyzed but yielded similar results (BF_01_ = 2.33 and 3.09, respectively).

**Figure 2.**
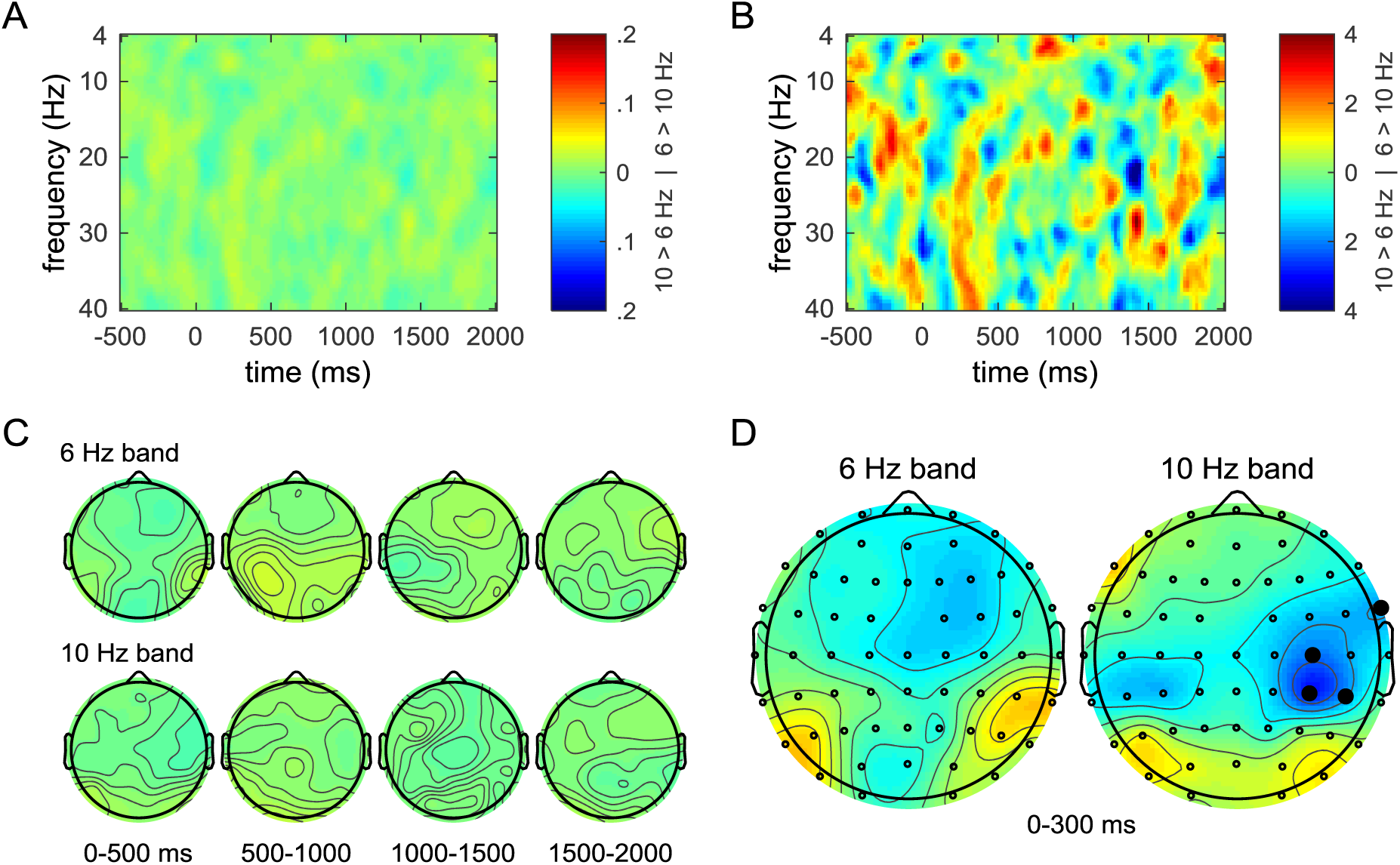
ITC during the indirect test phase. (A) Group-averaged time-frequency plots of ITC differences, collapsed over all electrodes. The color scale from Figure 1A is used to highlight the relatively limited range of differences during the indirect test phase compared to at encoding. (B) T-statistics associated with the differences from panel A. Note the restricted color scale compared to Figure 1B. As described in the text, none of the differences surpassed the cluster-corrected threshold. (C) Topographic maps of ITC differences for the 6-Hz (top row) and 10-Hz (bottom row) frequency bands, averaged into consecutive 500-ms latency periods. The color scale from panel A is used. (D) Topographic maps of t-statistics for ITC differences in the 6- and 10-Hz frequency bands, collapsed over the 0–300 ms period. Significant electrode-wise differences (p < .05, one-tailed) are designated by closed black circles, but the denoted clusters did not pass the corrected threshold. The color scale from panel B is used.

Evidence for ITC effects during the indirect test was further assessed according to its potential topographic distribution with two separate analyses. In the first analysis, reactivation effects were statistically tested in the 6-Hz and 10-Hz frequency bands at each electrode. Figure 2C displays the topographic maps of differences in these two bands averaged in 500-ms intervals across the recording epoch, and Figure 2D displays the resulting maps of t-values for the 0–300 ms period of interest. As shown, there were no significant effects for the 6-Hz band at any of the electrodes (maximum t_17_ = 1.48, p = .16, for electrode P8). In the 10-Hz band, there were significant effects at the FT8 (t_17_ = 1.85, p < .05), C4 (t_17_ = 2.28, p < .025), CP4 (t_17_ = 3.06, p < .005), and CP6 (t_17_ = 1.84, p < .05) electrodes, but the latter three clustered sites did not exceed the corrected threshold of 12 electrodes (corresponding to p < .05, one-tailed; the threshold for the 6-Hz band was 13 electrodes). Similar results were obtained from the analysis of the 0–500 ms interval, in which the largest cluster was only two significant electrodes (against a critical size of 12) for the 10-Hz band.

In the second analysis directed at identifying any evidence of reactivated ITC during the indirect test, both time and scalp location was allowed to vary simultaneously. All of the electrodes were included in this analysis, but the time dimension was simplified by averaging the data into 100-ms intervals throughout the post-stimulus period. Significant ITC differences for either the 6- or 10-Hz frequency bands could thus take the form of a combination of clustered time points and/or electrodes. The critical cluster sizes (corresponding to p < .05, one-tailed) determined for these analyses were 69 and 77 data points for the respective bands. However, the largest clusters were only 48 (for 6 Hz; p = .148) and 25 (for 10 Hz; p = .480) significant points in size.

#### 2.2.3. Direct test

As described in the Introduction, the direct test provides more ideal conditions to identify reactivation, as subjects were explicitly instructed to retrieve and respond according to the flicker information from encoding. Words only eliciting correct responses were included in these analyses, although given that performance was near chance for each subject, this does not indicate that frequency information was successfully retrieved. (As reported in the Supplemental Material, similar results were obtained when using all test items, regardless of accuracy.) Figures 3A and 3B display the ITC differences and corresponding t-values, respectively, collapsed over all electrodes across the recording epoch. No clusters of significant data points passed the permutation-based correction procedure. The largest clusters of significant effects were 37 points (p = .680) exhibiting higher ITC for 6-Hz trials and 49 points (p = .420) exhibiting higher ITC for 10-Hz trials, against respective critical sizes of 110 and 112 data points. Averaging the data from the 6- and 10-Hz bands over the 0–300 ms period again revealed no evidence of reactivation for the 6-Hz band (t_15_ = 1.07, p = .30; BF_01_ = 2.40) nor for the 10-Hz band (t_15_ = .82, p = .42; BF_01_ = 2.91). For the longer period of 0–500 ms, there was no evidence of reactivation from either band (6-Hz: t_15_ = 1.70, p = .11; BF_01_ = 1.21; 10-Hz: t_15_ = .04, p = .97; BF_01_ = 3.91).

**Figure 3.**
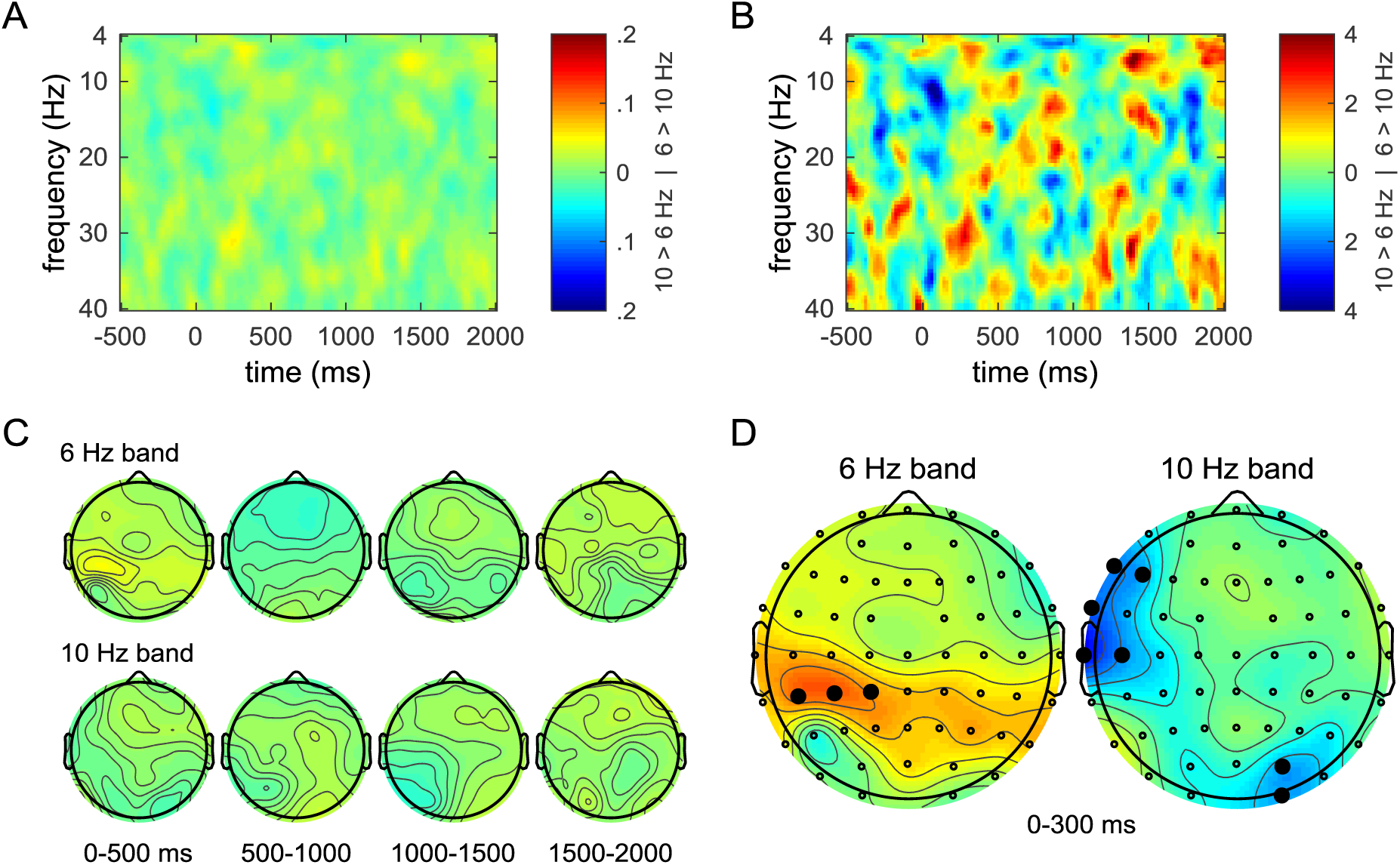
ITC during the direct test phase. (A) Group-averaged time-frequency plots of ITC differences, collapsed over all electrodes. The color scale from Figure 1A is used to highlight the limited range of differences during the direct test phase. (B) T-statistics associated with the differences shown in panel A. None of the differences surpassed the cluster-corrected threshold. (C) Topographic maps of ITC differences for the 6-Hz (top row) and 10-Hz (bottom row) frequency bands, averaged into consecutive 500-ms latency periods. The color scale from panel A is used. (D) Topographic maps of t-statistics for ITC differences in the 6- and 10-Hz frequency bands, collapsed over the 0–300 ms period. Significant electrode-wise differences (p < .05, one-tailed) are designated by closed black circles, but none of the denoted clusters passed the corrected thresholds. The color scale from panel B is used.

To assess topographic effects during the direct test, differences from the 6- and 10-Hz bands were tested at each electrode. Figure 3C displays the topographies of these differences across the recording epoch, whereas the resulting t-value maps from the 0–300 ms period are shown in Figure 3D. For the 6-Hz band, only three electrodes exhibited significant effects: CP5 (t_15_ = 2.22, p < .025), CP3 (t_15_ = 2.38, p < .025), and CP1 (t_15_ = 1.86, p < .05). For the 10-Hz band, there was a cluster of five significant electrodes over left fronto-central scalp (F7, F5, FT7, T7, and C5; t_15_ = 1.82 to 3.21, p = .044 to .003) and another cluster of two significant electrodes over right posterior scalp (PO4 and O2; t_15_ = 1.84 to 1.88, p = .042 to .040). Each of these effects failed to pass the cluster-corrected thresholds of 14 electrodes for both the 6- and 10-Hz bands. Similar analyses of data collapsed over the 0–500 ms period revealed that for the 6-Hz band, the left-lateralized cluster expanded to eight electrodes (FT7, T7, C5, C3, CP5, CP3, CP1, and CPz), and there was an additional cluster of three electrodes (P4, P6, and P8) over right posterior scalp. These effects again failed to surpass the appropriate cluster threshold of 17 electrodes. For the 10-Hz band over the 0–500 ms period, there were no significant electrodes. Finally, when the time dimension (100-ms intervals) was included in the topographic analyses, the largest clusters of time × electrode data points were 50 for the 6-Hz band (p = .155) and 18 for the 10-Hz band (p = .645). The respective cluster-wise thresholds for these analyses were 74 and 75 data points in size.

## 3. DISCUSSION

The current study used EEG to test the hypothesis that salient encoding information elicits neural reactivation at the time of retrieval, even when that information is presumably inaccessible to conscious awareness. A recent study by Wimber et al. (2012) reported a double dissociation of frequency-specific EEG effects during retrieval that corresponded to visual entrainment induced at encoding. Those findings were sought for replication here. As was shown by Wimber et al., statistically equivalent behavioral performance was observed in a recognition memory (indirect test) phase for words previously associated with two background flicker rates (6 and 10 Hz). These equivalent performance measures included overall hit rates, RTs corresponding to hits, and the proportions and RTs associated with high-confidence ratings. In a subsequent (direct) test phase that probed explicit retrieval of the frequency condition at encoding, subjects were also at chance in identifying the correct frequency and exhibited no RT differences across the prior frequency conditions. Consistent with the behavioral results of Wimber et al., these findings suggest that subjects did not, and were potentially unable to, consciously access the encoding information during retrieval.

EEG data were recorded during all phases of the current study – encoding and the indirect and direct tests – to investigate coherent (phase-locked) activation and reactivation due to visual entrainment. During encoding, the visual flicker elicited significant ITC differences at the stimulated frequencies as well as at higher frequencies (also see Herrmann, 2001). The encoding differences were evident across the scalp but additionally maximal at the posterior electrodes, as was anticipated due to the visual nature of the stimuli. Notably, the fact that the effect sizes for these across-scalp differences (Cohen’s d-values of 2.39 and 1.24 for 6 and 10 Hz, respectively) were as large as those obtained by Wimber et al. (2012; corresponding values of approximately 1.5 and 1, based on their Figure 2) suggests that the entrainment procedure was sufficient, all else being equal, to potentially elicit reactivation during the subsequent test phases. Despite the strong effects at encoding, however, no evidence of a frequency-specific dissociation was observed during either of the test phases. Specifically, such differences were not evident when: (a) collapsing across electrodes to increase statistical power, as was apparent during encoding; (b) testing frequencies other than those stimulated; (c) testing for effects occurring at any scalp location; and (d) conducting hypothesis-driven analyses based on the early time period (0–300 and 0–500 ms) of the possible effects (as in Wimber et al., 2012). Moreover, neither overall hits nor high-confidence judgments from the indirect test provided evidence of differences (see Supplemental Material), with high-confidence judgments potentially providing the better circumstances to observe such differences given the positive correlation between reactivation and confidence (Johnson et al., 2009, 2015). In sum, the ITC measures for both the indirect and direct memory tests were indistinguishable according to the prior frequency condition associated with the test words, constituting a failure to replicate the findings of Wimber et al. (2012).

Given the failure to replicate the critical finding of unconscious reactivation shown by Wimber et al. (2012), it is natural to speculate about possible differences between the studies that might account for the disparate results. As described in the Introduction, the current study set out to reproduce the previous one as closely as possible, based on the methodological details provided. These included specific aspects of the design, such as the numbers of stimuli comprising each phase and the timing of stimulus presentation, as well as the EEG recording and analysis procedures. Notably, the main difference between the studies was that the same subjects completed all of the test phases in the current study, whereas Wimber et al. enlisted an additional subject sample for the direct test; however, because this test was administered at the end of the session (and it was not referred to in the experimenter’s instructions until right before its start), it should have had no impact on the previous test phases. In addition, some aspects of the present results can be taken as confirmation that the methods were similar to those previously employed. First, as noted above, the differences observed at encoding were comparable in terms of effect sizes to those demonstrated by Wimber et al. This is an important finding since, based on fMRI studies (Johnson & Rugg, 2007; Johnson et al., 2009), the potential reactivation effects at retrieval would presumably be smaller than the activation effects at encoding. Second, overall behavioral accuracy in the indirect test phase of the current study was almost as high as that in the Wimber et al. study (d′ = 1.11 and 1.25, respectively), with the measures based only on high-confidence responses also being similar (d′ = 1.20 and d′ ≈ 1.27, respectively; note that the latter value is estimated from their Figure 1). Although this replication attempt appeared to include the key features of the previous study that make it analogous to other, successful investigations of reactivating encoding content at retrieval, there nevertheless could have been other subtle differences that were overlooked.

An additional difference between the Wimber et al. (2012) study and ours concerns the statistical correction procedures employed to control for multiple comparisons in the topographic analyses. Whereas Wimber et al. treated each electrode independently, thus counting the total number of electrodes across the scalp that exhibited significant effects, our procedure considered the correlations often observed among proximal electrodes (also see Maris & Oostenveld, 2007). We argue that the latter procedure is more appropriate given the smoothed nature of the effects over broad sections of the scalp (see panels C and D of Figures 2 and 3). However, employing the “independent” form of permutation-based correction with our indirect and direct test data resulted in critical thresholds ranging between 12 and 18 total electrodes for the combinations of different test phases, frequency bands (6 and 10 Hz), and latency windows (0–300 and 0–500 ms). The effect closest to approaching its corresponding threshold was for the 6-Hz band in the 0–500 ms window of the direct test, where a total of 11 electrodes displayed significant effects against a threshold of 15. Thus, the overall evidence for reactivation still failed to pass these alternative (and arguably, less ideal) correction procedures. Moreover, the fact that these independent thresholds were only slightly higher than those determined from the clustering procedure adds further support to the notion that the ITC measure exhibits natural dependencies between electrodes.

Some other studies that have investigated the effects of sensory manipulations on encoding-retrieval overlap are worth discussing in light of the current findings. In a seminal study in this area, Gratton et al. (1997, Experiment 4 in particular) briefly presented abstract line stimuli to the left or right of fixation at encoding and later had subjects make recognition-memory judgments for centrally-presented stimuli. Lateralized differences in the event-related potentials (ERPs) recorded during recognition corresponded to the side of presentation at encoding. Gratton et al. (1997) additionally showed in a subsequent experiment (Experiment 5) that subjects were at chance in identifying the side of original presentation. Fabiani et al. (2000) extended these findings by showing that such lateralized ERP differences distinguished between true and false recognition of words presented in the context of a Deese-Roediger-McDermott paradigm (Deese, 1959; Roediger & McDermott, 1995). Because both true and false memories were designated with the same (“old”) response, but only true recognition was associated with lateralized ERP effects, the authors inferred that the effects were outside of conscious awareness (for analogous fMRI results, see Slotnick & Schacter, 2004). Additionally, in a recent EEG study by Waldhauser et al. (2016), object stimuli presented to the left or right of fixation at encoding were associated with early (100–200 ms post-stimulus onset) lateralized oscillatory activity in the alpha/beta range (10–25 Hz) at the time of retrieval. Although there are several differences between the materials and methods of these studies and the current one, one difference stands out as potentially explaining the discrepancy in results: At some point, location information was made relevant to the task in all cases. That is, two of the studies involved explicit instructions to attend to the presentation locations of the stimuli (Gratton et al., 1997; Slotnick & Schacter, 2004; also see Slotnick & Schacter, 2010, and Thakral & Slotnick, 2014, 2015), another study used location to organize the words into semantically-related lists at the time of encoding (Fabiani et al., 2000), and the final study used a source memory task that required subjects to overtly respond to the location of encoding presentation. By comparison, the visual flicker in the current study was likely incidentally encoded, if at all, presumably leading to a decreased likelihood of being accessed again at retrieval.

Aside from any attempts to reconcile the current findings with those of previous studies, one can also speculate (with the help of hindsight, obviously) about other possible reasons for failing to demonstrate evidence of unconscious reactivation. One possibility is that the period shortly after word onset during the test phases might not be ideal for revealing coherent oscillatory activity (for evidence of early reactivation from other paradigms, see Jafarpour et al., 2014; Johnson et al., 2015; Waldhauser et al., 2016). Instead, this initial period might be dominated by the effects of word presentation, as was seemingly the case for the encoding data (see Figure 1A), where early ITC was negligible despite the visual stimulation being present. Even under the assumption that test words could quickly reactivate the associated encoding information, it seems likely that such an account would need to consider the time required for lexical access to occur, which is typically thought to be at least 100–200 ms (e.g., Bentin et al., 1999; Sereno & Rayner, 2003). Additionally, variability in the timing of such access, as would be expected given the specific characteristics of each word (e.g., length, frequency), might diminish the coherence of neural effects.

An alternative possibility is that, although the current study attempted to capitalize on the salience of visual entrainment (and consequently, its massive effects on the EEG), other factors might be driving cortical reinstatement. For example, the diagnosticity of encoding information to the recognition decision could carry more weight than salience. From this point of view, flicker stimulation differs from some of the manipulations that have typically been used to elicit reactivation, such as associating items with different encoding tasks (e.g., Kahn et al., 2004; Johnson & Rugg, 2007) or other classes of stimuli (e.g., Kuhl et al., 2011, 2013; Jafarpour et al., 2014). The associated information in those cases is presumably easier to access at the time of retrieval but is also often referred to explicitly during encoding to potentially modify the way in which items are processed. This delicate balance between providing potentially accessible information but encouraging subjects to avoid intentional retrieval highlights the difficulty of testing unconscious phenomena. Nonetheless, the consistent null results observed in the current study further contribute to the notion that boundary conditions exist around the involvement of reactivation in memory retrieval, and that one such boundary concerns the threshold of conscious awareness. These findings are thus compatible with neurobiological models positing that retrieval cues effectively able to engage hippocampally-mediated cortical reinstatement do so in service of providing conscious access to salient episodic information.

## 4. METHODS AND MATERIALS

### 4.1. Subjects

Twenty-five students from the University of Missouri (MU) participated in partial fulfillment of course credit. Informed consent was obtained in accordance with the MU Institutional Review Board. All subjects were right-handed, native-English speakers, with normal or corrected vision, and had no history of neurological disorders. The data from seven subjects were removed from all analyses, due to having excessive artifact in the EEG (three subjects) or inadequate behavioral performance (two subjects with hit minus false alarm differences < 0, and two subjects with too many missing test responses). The remaining sample of 18 subjects (6 females and 12 males) ranged in age from 18 to 22 years (M = 19). The data from two additional subjects were removed only from the analyses of the direct test phase, due to technical errors with EEG acquisition during that phase. (Including data from those two subjects in the behavioral analyses of the direct test did not change the pattern of results.)

### 4.2. Stimuli and design

A pool of 360 words was obtained from the MRC Psycholinguistic Database (Coltheart, 1981; Wilson, 1988; http://websites.psychology.uwa.edu.au/school/MRCDatabase/uwa_mrc.htm). Words were 4–9 letters long (M = 5.4), with written frequencies of 1–50 per million (M = 17.4; Kučera & Francis, 1967), and scores over 500 on scales of familiarity (M = 540), concreteness (M = 581), and imagability (M = 581). The words were randomly assigned to six lists of 60 words each that were rotated across the conditions across subjects. For each subject, words from four lists were presented during the encoding phase (one list assigned to each flicker frequency in each encoding block). The encoded words were presented again as old items during the corresponding *indirect test* block and in the final, *direct test* phase. Words from the remaining two lists served as new items during the indirect test. All words were shown in white uppercase Arial font (approximately 9 × 1 cm for the longest word) at the center of a black rectangular box (9.3 cm wide × 8.0 cm high) on the gray background of a 24-inch widescreen LCD monitor (cropped to 1024 × 768 resolution). During the encoding phase, the rectangular box flickered on and off the screen (i.e. between black and gray) at a frequency of either 6 or 10 Hz. The display was viewed at a distance of approximately 1 m. Stimulus presentation was controlled by the Cogent 2000 toolbox (http://www.vislab.ucl.ac.uk) in MATLAB (The MathWorks, Natick, MA).

### 4.3. Procedure

After being fitted with an electrode cap (lasting about 30 minutes), two cycles of an encoding block and an indirect test block were then completed, followed by a single direct test at the end of the session. Practice was administered before each block to ensure understanding of the procedures, and subject-paced breaks were provided at the midpoint of each block.

In each encoding block, 120 words were presented visually, one at a time, and subjects had to judge whether each word had an odd or even number of syllables. Words were presented for 2500 ms on a background rectangular box that flickered at either 6 or 10 Hz (60 trials of each condition per encoding phase). The two flicker conditions were randomly intermixed during each encoding phase. A white question mark followed word presentation and was displayed centrally for 1000 ms to signify that subjects should make their response. Subjects used their right index and middle fingers to indicate “odd” (“j” key) and “even” (“k” key) responses on the keyboard. A white fixation cross then appeared centrally for a random interval between 750 and 1250 ms until the start of the next trial.

Following the first encoding block, subjects completed a brief practice version of the indirect memory test and were further informed that they were being tested only on words from the immediately preceding block. Each indirect test block consisting of intermixed presentation of the 120 words from encoding and 60 new words. The words were presented against a black rectangular box for 2000 ms each. A question mark then prompted subjects to make their response and remained on the screen for 1500 ms. Subjects were instructed to indicate their confidence that each word was old or new using a 6-point scale: “sure old”, “probably old”, “maybe old”, “maybe new”, “probably new”, and “sure new”, respectively mapped to the “z”, “x”, “c”, “,”, “.”, and “/” keys. The keys were to be pressed with the index, middle, and ring fingers of each hand. A central fixation cross followed the response prompt and was displayed for a random interval between 750 and 1250 ms.

Following the second cycle of encoding and indirect test, the direct memory test was administered. The direct test consisted of the 240 words from the encoding blocks presented in a new random order. Words were presented on the black rectangular box for 2000 ms each and were followed by a question mark for an additional 1500 ms. While the question mark was displayed, subjects were instructed to indicate whether the rectangular background flickered at a “slow” (6 Hz) or “fast” rate (10 Hz) when the word was presented previously at encoding. The “j” and “k” keys were respectively used for these responses. A central fixation cross appeared for the randomly jittered interval between 750 and 1250 ms between trials.

### 4.4. EEG acquisition and analysis

EEG was continuously recorded during all phases of the experiment. Data were acquired with a BrainAmp Standard system (Brain Vision LLC; Durham, NC; http://www.brainvision.com) from 59 Ag/AgCl ring electrodes embedded in an elastic cap (Easycap, Herrsching, Germany; http://www.easycap.de). The electrode locations were based on the extended 10–20 system (Chatrian et al., 1985; Easycap montage 11) and included the following chains of sites: Fpz/1/2; AFz/3/4/7/8; Fz and F1 through F8; FC1 through FC6 and FT7/8; Cz, C1 through C6, and T7/8; CPz, CP1 through CP6, and TP7/8; Pz and P1 through P8; POz, PO3/4/7/8; and O1/2. Data were recorded with reference to an electrode placed at the FCz location, a ground electrode was placed at FT10, and additional electrodes were adhered to the mastoids (M1/2), below the left eye (IO1), and on the outer canthi (LO1/2). Electrode impedances were adjusted to below 5 kΩ before the experimental procedure began. Data were recorded at a sampling rate of 1 kHz and amplifier bandwidth of 0.01–100 Hz.

Offline processing of the EEG data was implemented with the EEGLAB (Delorme & Makeig, 2004; http://sccn.ucsd.edu/eeglab/) toolbox in MATLAB. The data were downsampled to 200 Hz, high-pass filtered at .05 Hz, re-referenced to the mastoid average, epoched from -1000 to 2495 ms relative to each word onset, and baseline-corrected to the pre-stimulus interval. Independent components analysis (ICA) was used to identify artifacts (e.g., eye movements, blinks, and muscle activity) that were then manually rejected based on their scalp topographies and power spectra (see Jung et al., 2000). The ICA procedure was applied to the data concatenated from all of the experimental phases to minimize any bias of the removal of components for one phase (or condition) versus another. Finally, the data were then low-pass filtered at 50 Hz.

The data were next time-frequency transformed into 73 frequency bands (4 to 40 Hz in .5-Hz intervals) and 125 time points (20-ms bins across epochs of -500 to 2000 ms) with a Morlet-wavelet procedure (3 cycles at the lowest frequency, linearly increasing to 15 cycles at the highest frequency; also see Delorme & Makeig, 2004). *Inter-trial coherence* (ITC; sometimes called *phase-locking factor*, as in Tallon-Baudry et al., 1996) was calculated separately for each electrode and each condition of interest in all of the experimental phases. ITC varies between 0 and 1, with larger values indicating more synchronous activity across trials and smaller values corresponding to desynchronized (random) activity. Given that ITC can be positively biased by the number of trials included in the analysis, particularly when ITC is low (Vinck et al., 2010), we equated the trial numbers per condition via random selection. For the indirect and direct test phases, respectively, averages of 86 (SD = 15; minimum = 53) and 53 (SD = 6; minimum = 40) trials per condition contributed to the analyses. Finally, due to the potential non-normality of the ITC measures, Wilcoxon signed-rank tests (also see Wimber et al., 2012) were employed in addition to standard t-tests. Note that the conclusions drawn with the two measures were similar; the t-test results are thus reported in the main text, while the results from signed-rank tests are relegated to the Supplemental Material. The report of t-tests also allowed for maintaining consistency with other results based on Bayes-factor t-tests and ANOVAs (see below).

Permutation-based analyses were used to control the family-wise error rate (FWER) due to the multiple comparisons across time, frequency bands, and electrodes (see Maris & Oostenveld, 2007). Neighboring electrodes were designated according to the appropriate montage from the FieldTrip toolbox (“Easycap M1”; Oostenveld et al., 2011; http://fieldtrip.fcdonders.nl). All other data points (i.e., frequency bands or time points) were treated as neighbors according to 18-edge connectivity (based on the Statistical Parametric Mapping toolbox; http://www.fil.ion.ucl.ac.uk/spm). For a given analysis, null distributions were created by randomly shuffling the condition labels (i.e., 6 versus 10 Hz) for each subject 1000 times and then, for each shuffling, testing the significance of each data point at the group level. The maximum-sized clusters for each shuffling were sorted to determine the critical cluster size (*k*) across permutations. Results in the time × frequency domain were thresholded at p < .025 in each direction (6- > 10-Hz trials, and vice versa), given that both directions of effect occupied the same search space. Results restricted to one frequency band, such as was the case for topographic maps and topography × time data, were thresholded at p < .05 for only the predicted direction of effect (i.e., ITC in the 6-Hz band was expected to be higher for 6- than 10-Hz trials, and vice versa). The p-values reported for analyses controlled with these methods correspond to the proportion of permutations (out of 1000) that gave rise to larger maximum-sized clusters than the actual results.

Finally, to indicate the relative evidence for the null and alternative hypotheses, Bayes factors (BFs) were computed for each statistical test using the BayesFactor package (http://cran.r-project.org/web/packages/BayesFactor/index.html; Rouder et al., 2009, 2012, 2016) for R (http://www.r-project.org). Each calculation assumed a JZS prior (Jeffreys, 1961; Zellner & Siow, 1980) and a pre-determined scaling factor (*r*) of .707 (based on the default setting in the BayesFactor package). BFs favoring the null and alternative hypotheses are respectively denoted as BF_01_ and BF_10_ and represent an odds ratio of the relative support. For example, a BF_01_ of 3 represents the data being three times more likely to have occurred under the null hypothesis than under the alternative hypothesis. BFs can be summarized with descriptive thresholds such as “anecdotal” (≥ 1), “moderate” (≥ 3), and “strong” (≥ 10; Jeffreys, 1961; Nuijten et al., 2015).

## SUPPLEMENTAL MATERIAL

**Figure S1.**
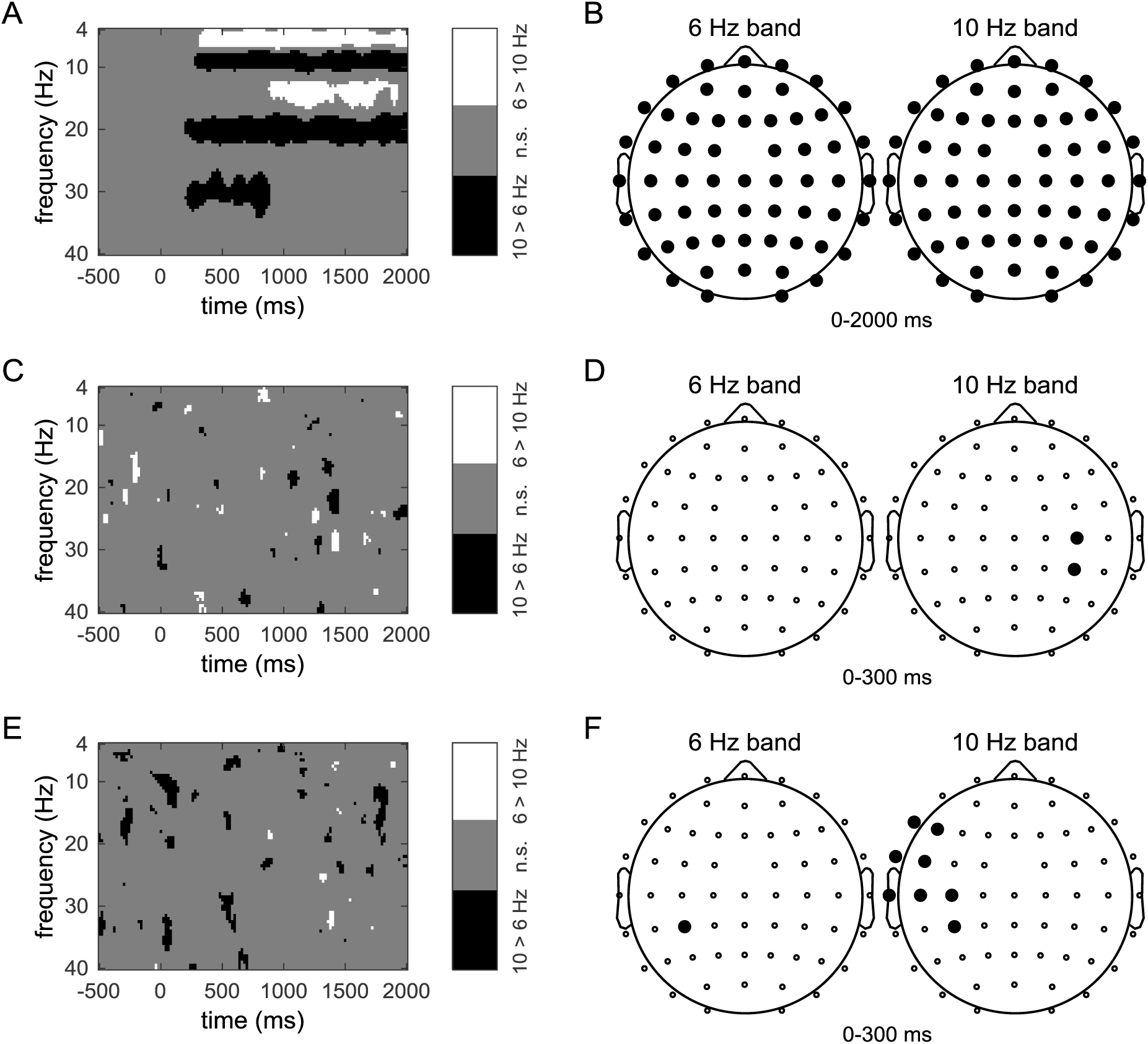
ITC differences during each experimental phase, based on Wilcoxon sign-rank tests. (A) Significant clusters of ITC differences during the encoding phase, across the analyzed frequency spectrum and trial epoch. Data are collapsed over all electrodes. White denotes higher ITC for 6-Hz trials, black denotes higher ITC for 10-Hz trials, and gray denotes non-significant data points. Data points that reached significance on their own, but not at the cluster level, are not displayed. (B) Montages of electrodes exhibiting significant ITC differences during the 0–2000 ms period of encoding phase for the 6-Hz and 10-Hz frequency bands. Significant differences were evident at every electrode, as indicated by the closed black circles. (C) Time-frequency plots of significant ITC differences during the indirect test phase, collapsed over all electrodes. None of the clusters surpassed the cluster-corrected threshold. (D) Montages of electrodes exhibiting significant ITC differences (denoted by closed black circles) during the 0–300 ms period of the indirect test phase for the 6-Hz and 10-Hz frequency bands. The denoted cluster did not pass the corrected threshold. (E) Time-frequency plots of significant ITC differences during the direct test phase, collapsed over all electrodes. None of the clusters surpassed the cluster-corrected threshold. (F) Montages of electrodes exhibiting significant ITC differences (denoted by closed black circles) during the 0–300 ms period of the direct test phase for the 6-Hz and 10-Hz frequency bands. The denoted clusters did not pass the corrected threshold.

**Figure S2.**
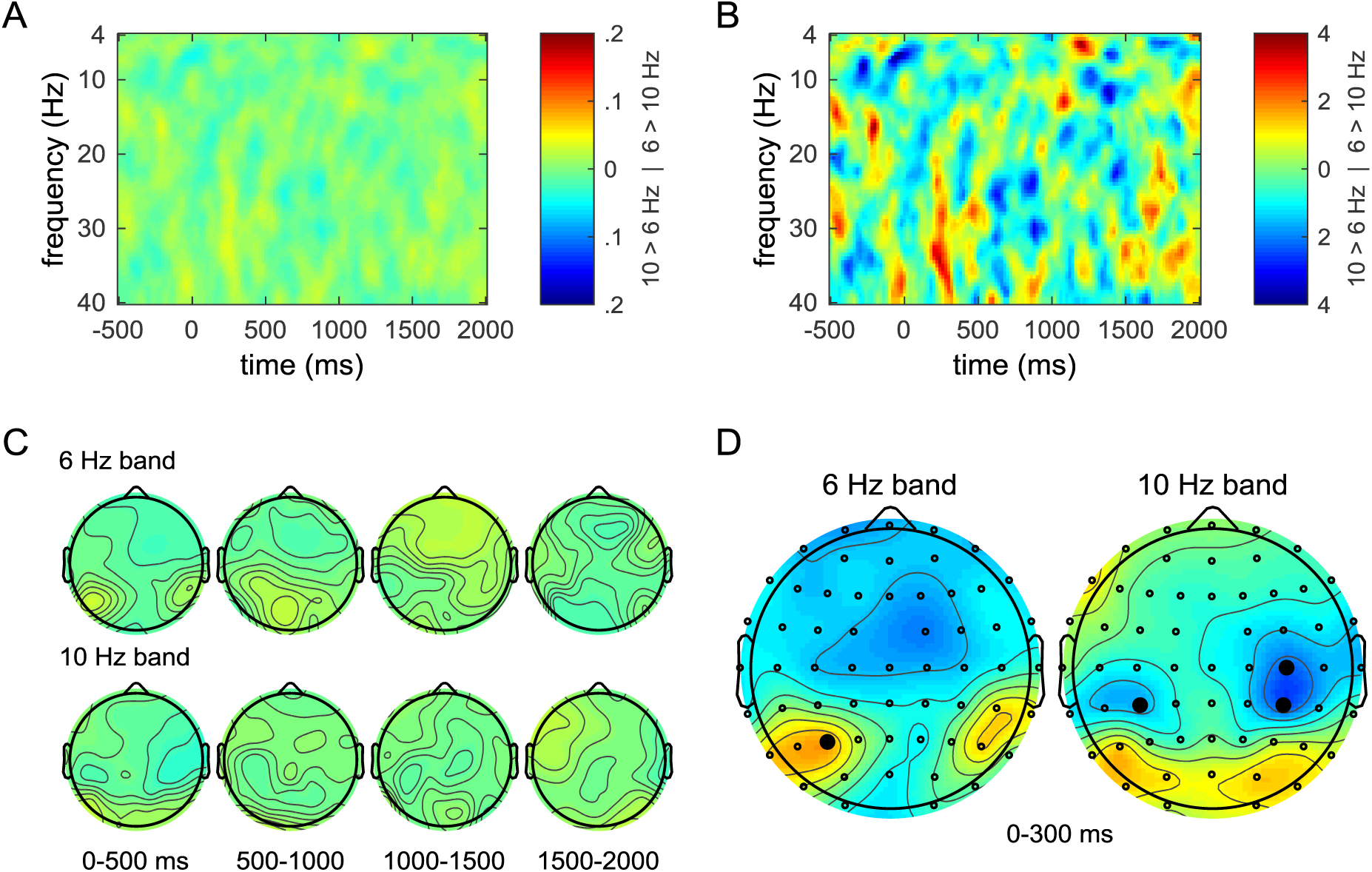
ITC during the indirect test phase, based only on high-confidence responses. Compare with Figure 2 from the main text, which is based on all correct old responses (hits). (A) Group time-frequency plots of ITC differences, collapsed over all electrodes. Warm colors indicate higher ITC for 6-Hz trials; cool colors indicate higher ITC for 10-Hz trials. (B) T-statistics associated with the differences from panel A. None of the differences surpassed the cluster-corrected threshold. (C) Topographic maps of ITC differences for the 6-Hz (top row) and 10-Hz (bottom row) frequency bands, averaged into consecutive 500-ms latency periods. The color scale from panel A is used. (D) Topographic maps of t-statistics for ITC differences in the 6- and 10-Hz frequency bands, collapsed over the 0–300 ms period. Significant electrode-wise differences are designated by closed black circles, but the denoted effects did not pass the cluster-corrected threshold. The color scale from panel B is used.

**Figure S3.**
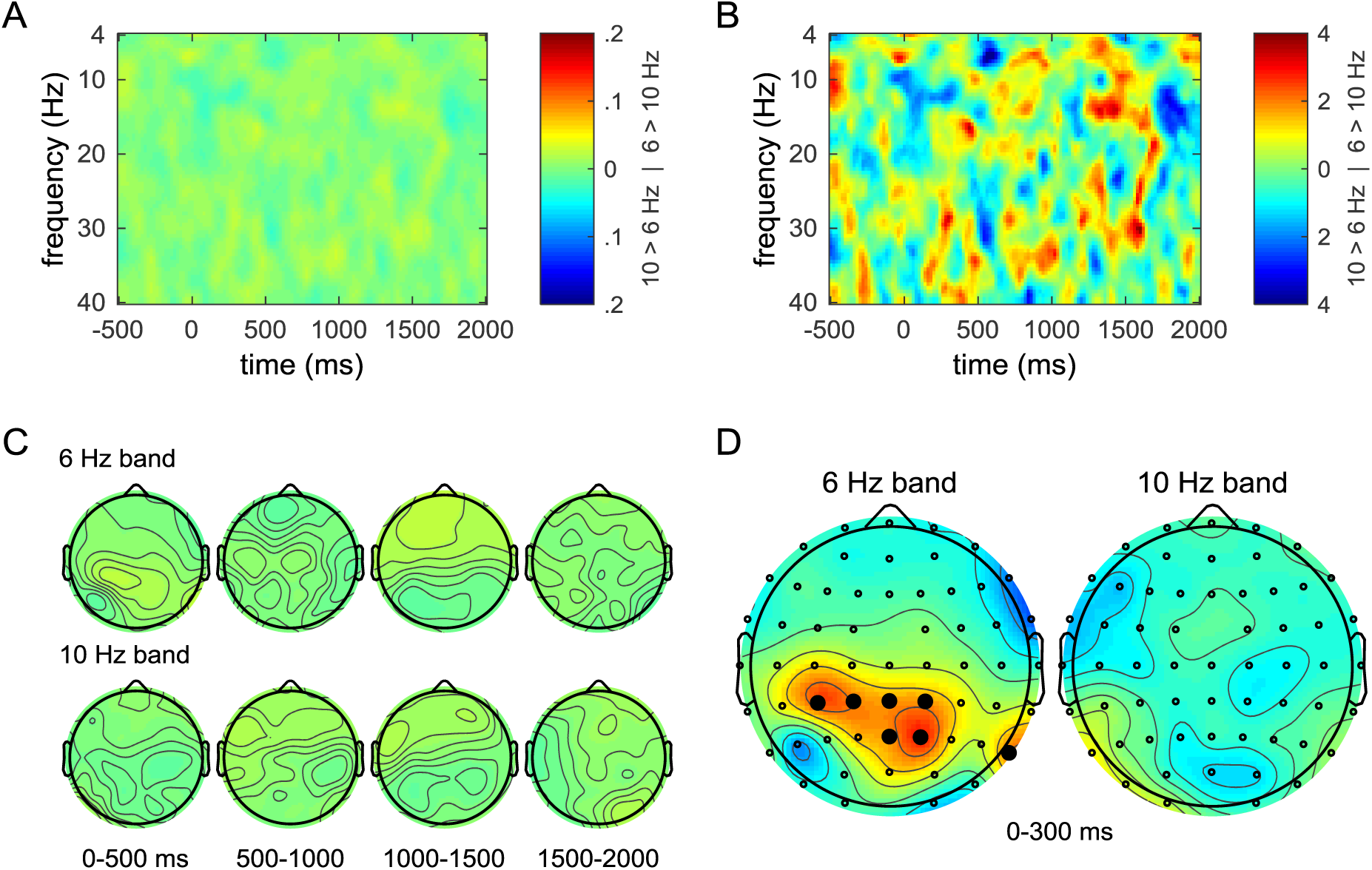
ITC during the direct test phase, based on all responses (regardless of accuracy) Compare with Figure 3 from the main text, which is based only on the correct responses to prior frequency condition. (A) Group time-frequency plots of ITC differences, collapsed over all electrodes. Warm colors indicate higher ITC for 6-Hz trials; cool colors indicate higher ITC for 10-Hz trials. (B) T-statistics associated with the differences from panel A. None of the differences surpassed the cluster-corrected threshold. (C) Topographic maps of ITC differences for the 6-Hz (top row) and 10-Hz (bottom row) frequency bands, averaged into consecutive 500-ms latency periods. The color scale from panel A is used. (D) Topographic maps of t-statistics for ITC differences in the 6- and 10-Hz frequency bands, collapsed over the 0–300 ms period. Significant electrode-wise differences are designated by closed black circles, but the denoted effects did not pass the cluster-corrected threshold. The color scale from panel B is used.

**Figure S4.**
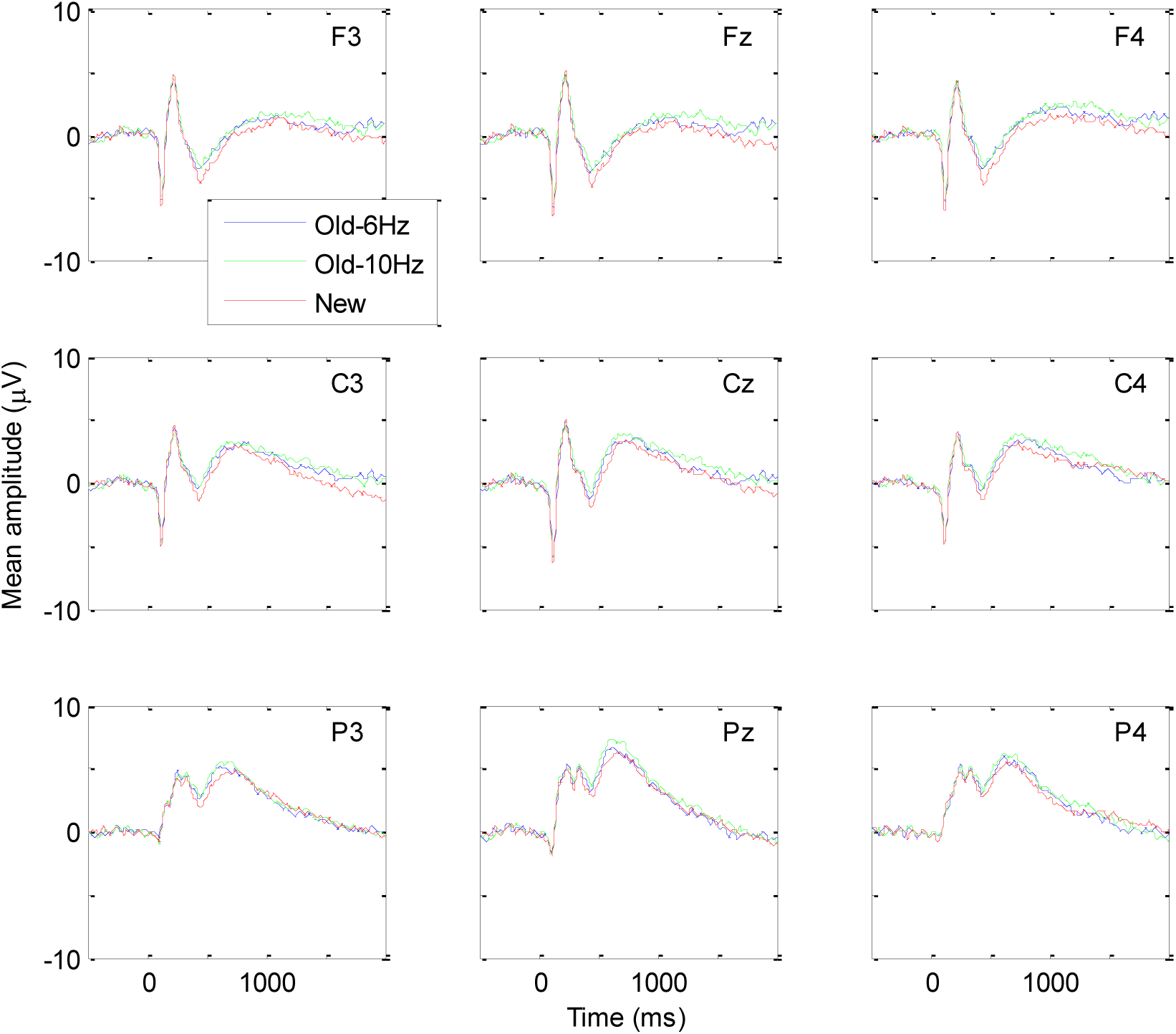
Grand-average ERPs for hits and correct rejections during the indirect test phase, collapsed across confidence level. ERPs elicited during the indirect test phase by old items associated with the 6- (blue) and 10-Hz (green) conditions and new items (red). Data correspond only to correct responses and are collapsed according to the three levels of confidence (“sure”, “probably”, and “maybe”). The electrode labels are indicated in the upper right corner of each plot.

**Figure S5.**
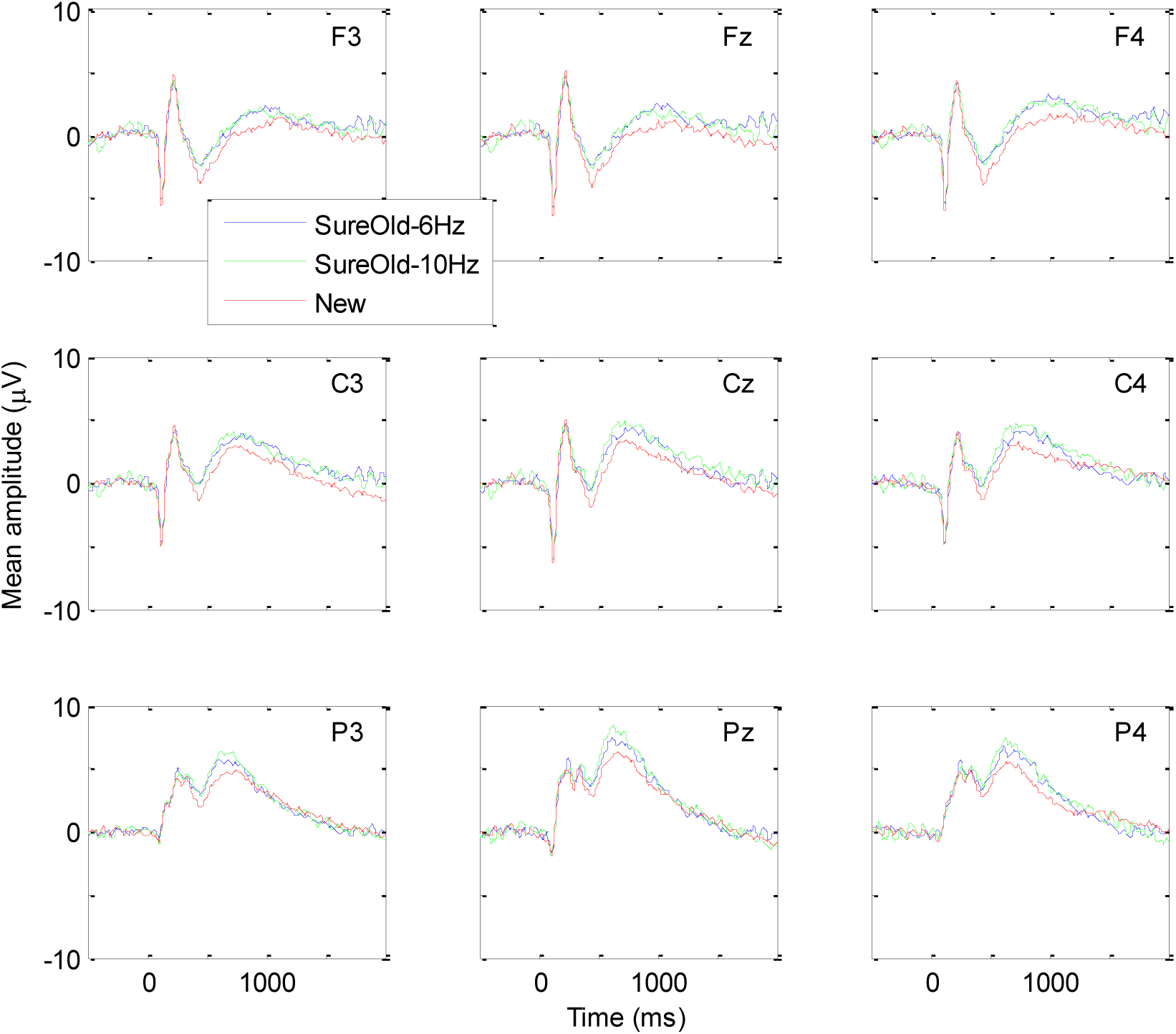
Grand-average ERPs for sure-old hits and correct rejections during the indirect test phase. ERPs elicited during the indirect test phase by old items associated with the 6- (blue) and 10-Hz (green) conditions and new items (red). The waveforms for old items correspond only to correct responses that were designated with “sure old” responses; correct responses to new items are collapsed across confidence levels (as in Figure S4). The electrode labels are indicated in the upper right corner of each plot.

## Footnotes

1 The term “incidental” is sometimes used in this context to refer to an effect that is irrelevant to the goals of the subject. In our opinion, however, the term can imply that information is successfully retrieved, even though it might be done in an involuntary or automatic manner (also see Kuhl et al., 2013, for discussion). As Wimber et al. (2012) demonstrated, and we show in the current study, there is no behavioral evidence for conscious retrieval of the flicker information. Thus, to minimize confusion, we use the term “indirect” to refer to the nature of the memory task rather than its outcome.

